# Exploring variations in lipids among drug-resistant and sensitive *Mycobacterium tuberculosis* by Thin layer chromatography and mass spectrometry

**DOI:** 10.1101/2024.03.22.586225

**Authors:** Kavitha Kumar, Prashant Giribhattanavar, B. K. Chandrasekhar Sagar, Shripad A. Patil

## Abstract

*Mycobacterium tuberculosis (M.tb)* lipids are important in the host–pathogen interplay, variation in the lipids organization of cell wall can act as an adaptive response. Specific cell wall structures can possibly result in suboptimal intracellular concentrations of anti-TB drugs, which favors the acquisition of drug resistance. Therefore, lipids from *M.tb* (drug resistant and sensitive) were analyzed by 2D-thin layer chromatography and mass spectrometry. GraphPad Prism was used to perform Mann Whitney-U test to determine the statistical significance. Difference observed for total lipid content among different resistant isolates was insignificant. However, increase in phospholipids was identified in multi-drug resistant (MDR) isolate compared to sensitive isolate. Isoniazid, streptomycin-isoniazid, and isoniazid-ethambutol resistant isolates showed increased alpha-mycolic acids. MDR isolate showed a marginal decrease in alpha- and keto-form. Mycolipenic acid was seen only in sensitive isolate, and mycosanoic acids were observed in all the resistant isolates. Among the resistant isolates, there was an insignificant increase in the total phthiocerol dimycocerosates and sulfolipids. Drug resistance was associated with compositional imbalance of lipids. However, investigations to determine whether the changes notices are induced by the drugs is to be explored, which could give an insight into the drug resistant organisms pathogenesis.

## 1. Introduction

The outcome of tuberculous meningitis (TBM) with drug-resistant *Mycobacterium tuberculosis (M.tb)* is often poor and associated with exceptionally high mortality. Globally in 2015, an estimated 3.9% and 21% of new cases and previously treated cases had multidrug resistant-tuberculosis (MDR-TB) respectively. In India, MDR-TB is among 2.5% and 16% of new cases and previously treated cases respectively. Russian Federation, India and China are with most (45% of the global total) numbers of MDR cases (1). Drug resistant form of TBM is difficult to diagnose and treat as the targeted regimens are still unexplored (2).

The *M.tb* cell wall grows in snake-like cords and the most complex of all bacteria. The most distinctive feature is that lipids with mycolic acids (MA) occupies 60% of the weight. Many drugs used to combat mycobacteria target the mycolyl-arabinogalactan-peptidoglycan (mAGP) complex which forms major part of the cell wall. Additionally phthiocerol dimycocerosates (PDIM), glycopeptidolipids (GPL), menaquinones (MQ) and glycosylated phenolpthiocerols lipids intercalate to the MA layer and form the cell wall outer region (3) Research over the past has implicated the importance of *M.tb* cell wall for the host-pathogen cross-talk and also for pathogen. As the drug-resistant strains are increasing globally, cell wall impact with respect to resistant trait becomes vital.

The study objective was to investigate the drug-resistant *M.tb* lipids and gain further understanding of first line drug resistance in *M.tb*. Hence, lipids from drug-resistant and drug sensitive Mycobacteria was compared to assess the compositional changes.

## 2. Materials and Methods

### 2.1 Samples

*M.tb* organisms (n=42) were isolated from cerebrospinal fluid (CSF) of TBM patients admitted at National Institute of Mental Health and Neurosciences during 2013-2014. The study was approved by the Institutional Ethics Committee. Isolates were analyzed for their susceptibilities to first-line drugs: isoniazid (I), rifampicin (R), streptomycin (S), and ethambutol (E) by Mycobacteria Growth Indicator Tube (MGIT) method as per manufacturer’s instructions (4). Lipids was analyzed among sensitive and different resistant categories that includes mono-drug, bi-drug, poly-drug and multidrug resistance. H37Rv was used as the reference standard for the analysis. All the experiments were conducted in biosafety level 3 (BSL3) lab.

### 2.2 Extraction of Lipids

Modified Chandramauli’s method was used to extract lipids as given by Singh et al., (5) with slight modification. Briefly, 1g of *M.tb* cells grown on solid LJ media was mixed with Chloroform: methanol (2:1) mixture (6ml), and kept on a stirrer for 12 hrs. The suspension was centrifuged, and organic layer was collected. The step was repeated for the residue with the 4ml of chloroform: methanol (2:1). The pooled mixture was washed with 3ml of 0.29% NaCl, organic phase was collected and kept for drying. The dry weight of the residue was recorded.

### 2.3 Lipid Analysis by 2D Thin Layer Chromatography (TLC)

Extracted lipid (dried organic mixture) was solubilized in chloroform and 4µg was resolved by 2D TLC on aluminum-backed silica gel plates (Merck, India) by spotting the extract using microcapillary pipettes as described Pirson et al., (6). Separate solvent system was used for analysis of non-polar and polar lipids (**Table 1**).

**Table 1:**
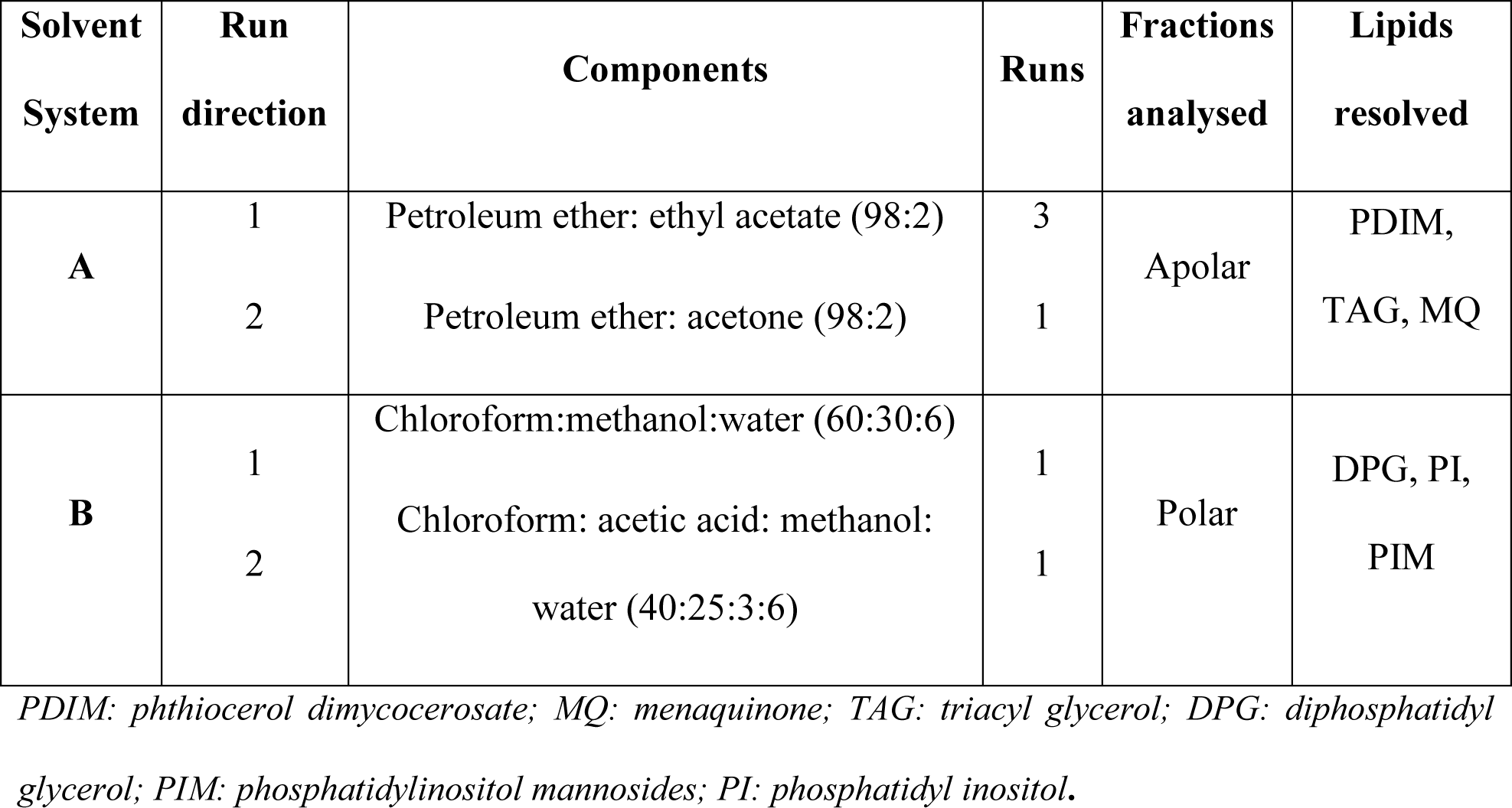
Solvent system for 2D-Thin Layer Chromatography analysis of lipids.

Staining was using 5% phosphomolybdic acid (PMA) solution (SRL chemicals, India) in 95% ethanol. Stains were sprayed and charred in hot air oven.

Individual lipids were identified by comparing with published TLC analysis (6). The image of TLC plates was used for quantitative assessment of individual spots in the JustTLC software (Sweday). Each spot was marked to cover the entire area of the spot on the TLC plate, measured relative density which was calculated as a percentage of the total plate and statistical significance was determined.

### 2.4 Lipids Analysis by Mass spectrometry

Lipids extracted from the selected clinical isolates (Resistant to I, MDR (resistant to IR), SI resistant, IE resistant and sensitive isolates (susceptible to SIRE) was subjected to Mass spectrometry.

Briefly, extracted lipid sample was solubilized in dihydroxybenzoic acid (DHB) matrix. The samples were spotted on MALDI target plate and analysed on Bruker UltrafleXtreme mass spectrometer (Bruker Daltonics, Bremen, Germany) with positive voltage polarity mode. Data was collected as profiled spectra over a mass range of 300 to 3,500 Da using Flex Analysis 3.1 software. Mass-to-charge ratio (m/z) values obtained were matched from *M.tb* lipidome present in Mass Spectrometry-based Lipid(ome) Analyzer and Molecular Platform (MS-LAMP) software with 0.5 window. The lipids matched was used to compare between the sensitive and different categories of resistant isolates.

### 2.5 Statistical analysis

GraphPad online software was used for data analysis. Statistical significance was determined at *P* <0.05. Relative density of lipids from sensitive and resistant isolates were analyzed using the Mann-Whitney U test.

## 3. Results

### 3.1 Lipids Analysis by 2D TLC

Total lipid content was analyzed and relative densities of lipid fractions from sensitive and resistant isolates revealed changes in the profile **(Table 2)**. The variation in dry weight was statistically insignificant. Lipid fraction analysis by solvent system A showed non-polar lipids (**Figure 1**) and system B enabled the identification of polar lipids (**Figure 2**).

**Figure 1:**
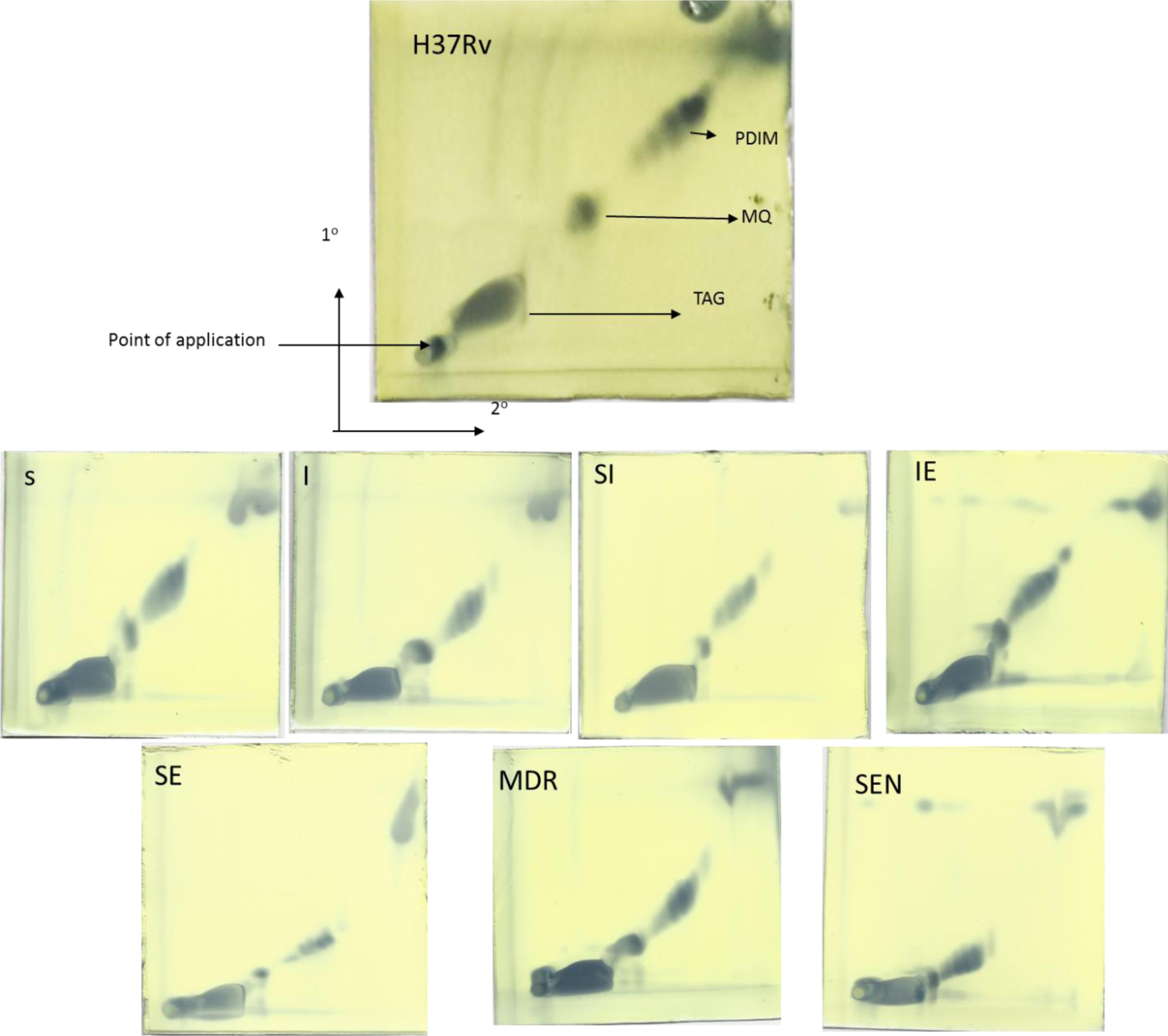
Representative 2D-Thin layer chromatogram of extracted lipids showing the presence of non-polar lipids. PDIM: phthiocerol dimycocerosate; MQ: menaquinone; TAG: triacyl glycerol; H37Rv: Standard laboratory strain; S: Streptomycin; I: Isoniazid; R: Rifampicin; E: Ethambutol; MDR: Multidrug resistant; SEN: Sensitive; 1°: first run direction of solvent system; 2°: second run direction of solvent system.

**Figure 2:**
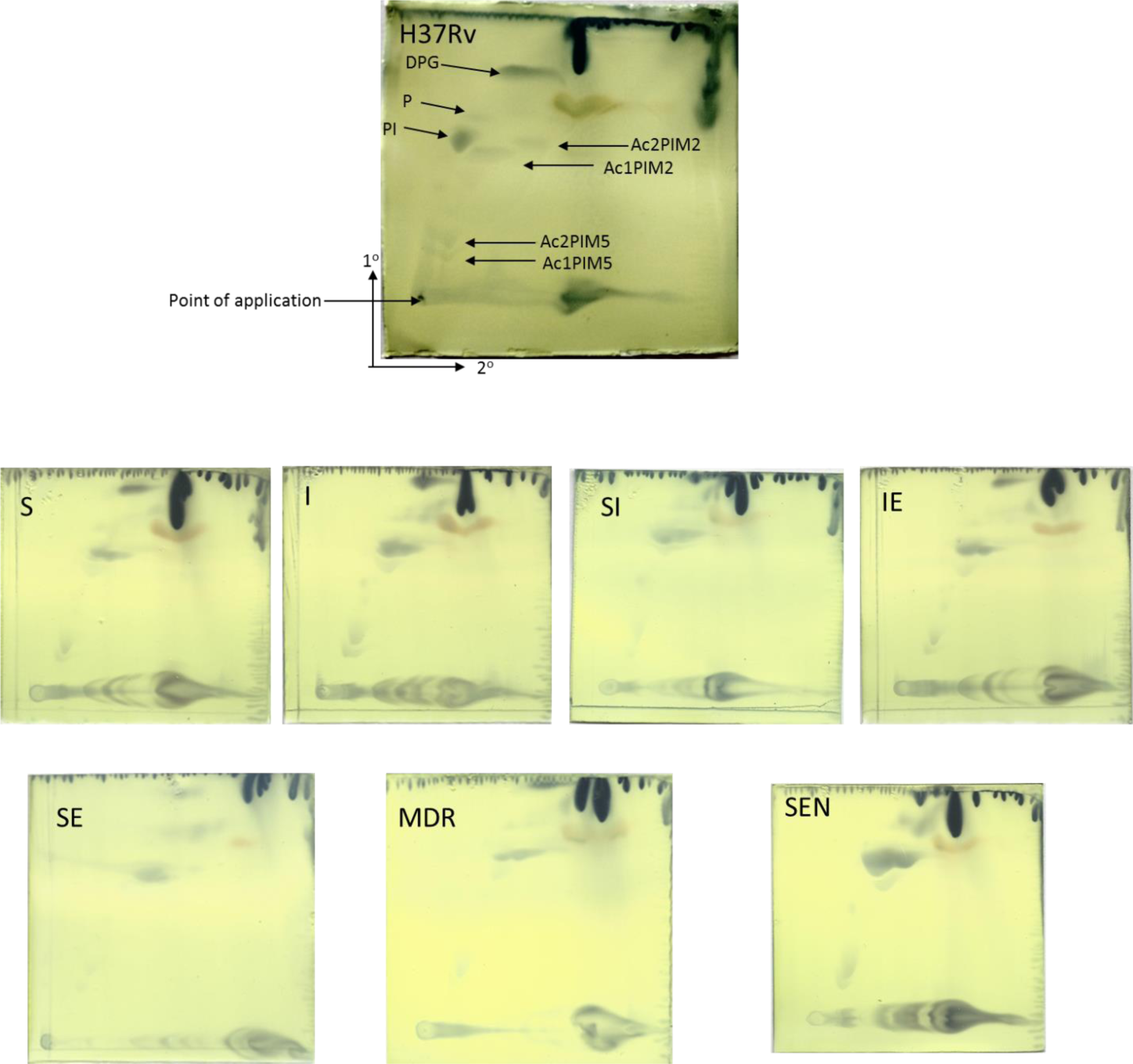
Representative 2D-Thin layer chromatogram of extracted lipids showing the presence of polar lipids. DPG: diphosphatidyl glycerol; PIM: phosphatidylinositol mannosides (integers denote number of mannoside or acyl groups); PI: phosphatidyl inositol; P: phospholipid; H37Rv: Standard laboratory strain; S: Streptomycin; I: Isoniazid; R: Rifampicin; E: Ethambutol; MDR: Multidrug resistant; SEN: Sensitive; 1°: first run direction of solvent system; 2°: second run direction of solvent system.

**Table 2:**
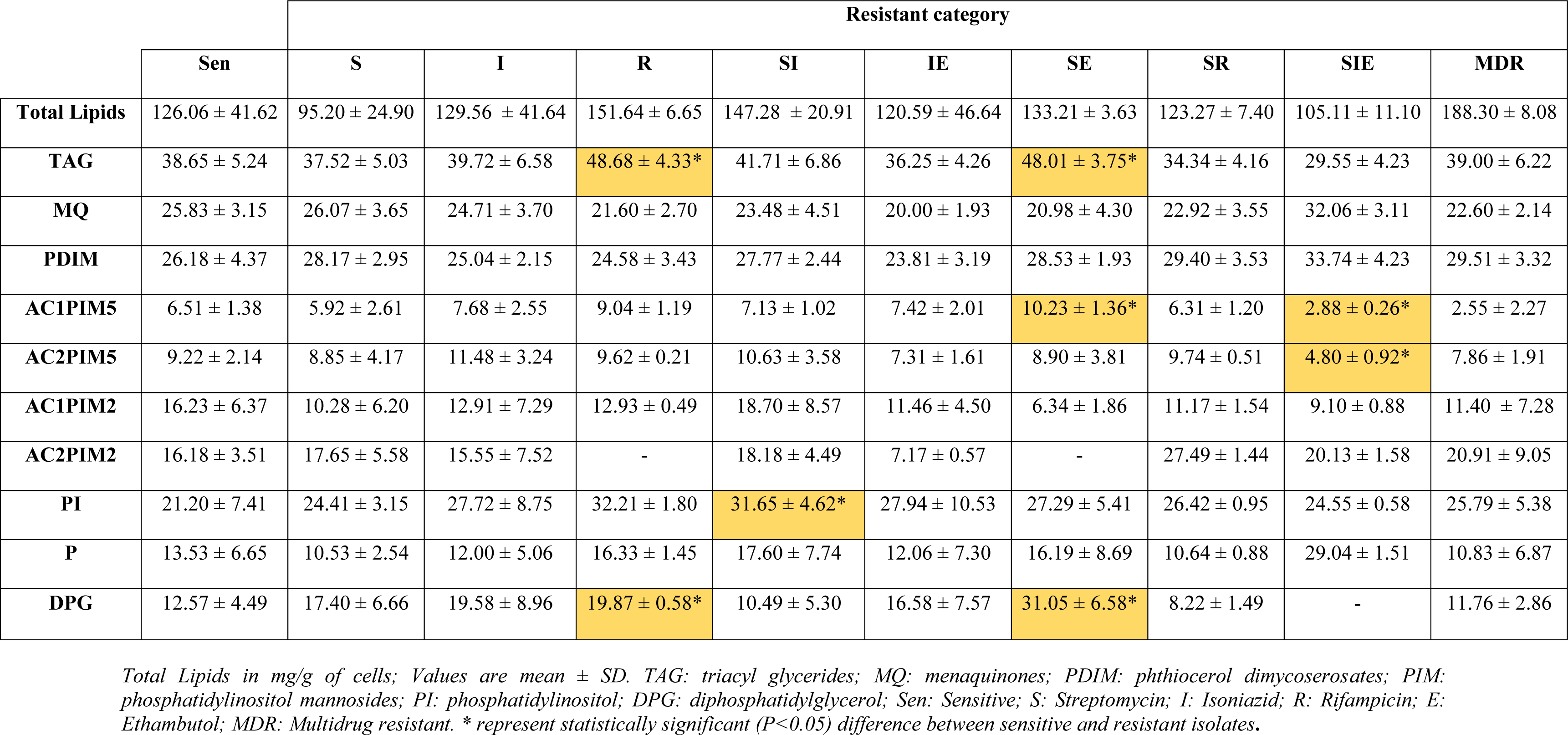
Comparison of relative density of different lipids among sensitive and resistant *M.tb* clinical isolates.

The analysis of individual fractions revealed a significant difference in few of the lipids analysed **(Table 2)**. Among non-polar lipids, Triacyl glycerides (TAG) were higher significantly (*P*<0.036) in rifampicin (R) resistant and streptomycin-ethambutol (SE) resistant isolates compared to sensitive isolate. Menaquinones (MQ) and phthiocerol dimycocerosate (PDIM) showed differences but it was statistically insignificant **(Table 2)**.

Among polar lipids, tri-acylated phospho-*myo*-inositol pentamannosides (AC1PIM5) was significant statistically (*P*<0.036) in streptomycin-ethambutol (SE) resistant isolate and streptomycin-isoniazid-ethambutol (SIE) resistant isolate and tetra-acylated phospho-*myo*-inositol pentamannosides (AC2PIM5) was significant statistically (*P*<0.036) in streptomycin-isoniazid-ethambutol (SIE) resistant isolate. Phosphatidyl inositol (PI) was statistically significant (*P*<0.032) in streptomycin-isoniazid (SI) resistant isolates. Diphosphatidylglycerol (DPG) was significantly higher (*P*<0.036) in rifampicin (R) resistant and streptomycin-ethambutol (SE) resistant isolates. **(Table 2)**.

### 3.2 Lipids analysis and comparison by Mass Spectrometry

The individual m/z values of the extracted lipids from one sensitive isolate and 4 resistant isolates was interpreted using the MS-LAMP software that classifies the lipids as 6 categories - fatty acyls (FA), glycerophospholipids (GP), glycerolipids (GL), polyketides (PK), prenol lipids (PR), and saccharolipids (SL).

There was variation in the lipids identified among the clinical isolates between sensitive and resistant isolates. The maximum number of lipids were obtained among MDR isolate (n=184) and the least was among SI resistant (n=158). FA was abundant in SI resistant isolate (23.42%), GL was abundant in sensitive isolate, and more number of lipids among all the 6 groups was found in GP, with more abundance in MDR isolate. PK, PR and SL were very few, among that PK and PR was more in IE category and SL was least among I resistant isolate (0.61%) (**Figure 3).**

**Figure 3:**
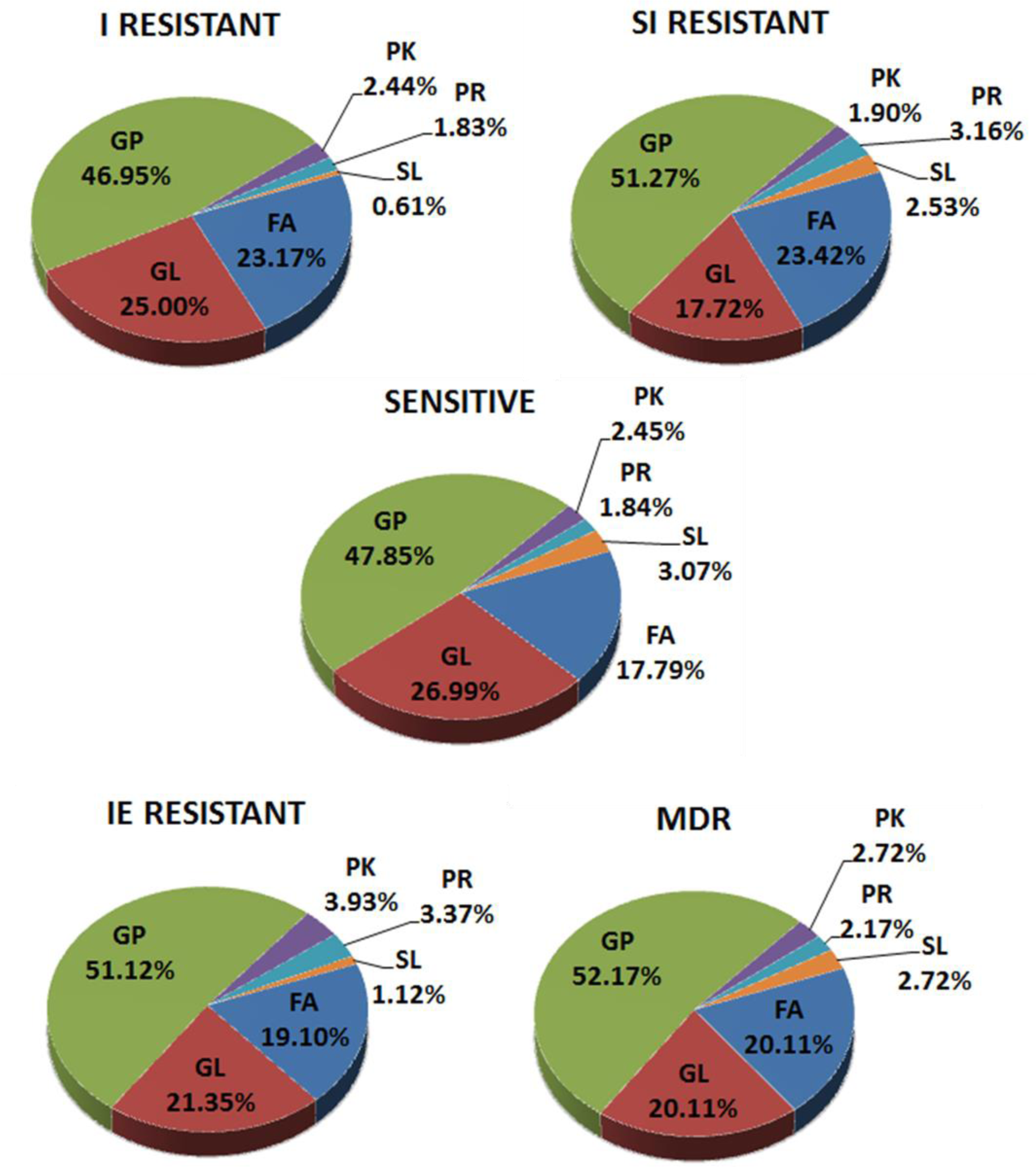
Pie chart of all the 5 isolates showing the 6 groups of lipids as obtained by MS-LAMP. Sen: Sensitive; S: Streptomycin; I: Isoniazid; R: Rifampicin; E: Ethambutol; MDR: Multidrug resistance; FA: Fatty Acyls; GL: Glycerolipids; GP: Glycerophospholipids; PK: Polyketides; PR: Prenol Lipids; SL: Saccharolipids.

Among the different categories of lipids, there are different sub-class of lipids. The FA category consisted of branched fatty acids that include mycocerosic acid, mycolipanolic acid, mycolipenic acid, mycosanoic acid, pthioceranic acid, hydroxypthioceranic acid etc; mycolates such as α-mycolic acids, methoxy mycolic acids, keto-mycolic acids, GMM, TMM and TDM; and fatty esters such as DIMA and DIMB along with their glycosylated counterparts were seen **(Table 3)**.

**Table 3:** Summary of different categories of lipids with their sub-class lipids found among the sensitive and resistant isolates of *M.tb*.

| LIPIDS |  | Sensitive isolate | Resistant isolates |  |  |  |
| --- | --- | --- | --- | --- | --- | --- |
| Categories | Sub-class |  | I | SI | IE | MDR |
| FA | Branched Fatty Acids | 14 | 14 | 10 | 11 | 13 |
|  | Mycolates | 11 | 20 | 21 | 18 | 17 |
|  | Fatty Esters | 4 | 4 | 6 | 5 | 7 |
| GL | Monoacylglycerols | 6 | 8 | 4 | 4 | 6 |
|  | Diacylglycerols | 18 | 16 | 9 | 14 | 12 |
|  | Triacylglycerols | 20 | 17 | 15 | 20 | 19 |
| GP | Glycerophosphoethanolamines (PEs) | 10 | 13 | 11 | 12 | 11 |
|  | Glycerophosphoglycerols (PGs) | 17 | 14 | 12 | 13 | 11 |
|  | Glycerophosphoinositols (PIs) | 9 | 12 | 13 | 15 | 12 |
|  | Glycerophosphoglycerophosphoglycerols (CLs) | 10 | 9 | 12 | 12 | 15 |
|  | Glycerophosphoinositolglycans | 32 | 29 | 33 | 39 | 47 |
| PK | Linear | 2 | 1 | 0 | 1 | 0 |
|  | Hybrid | 2 | 3 | 3 | 6 | 5 |
| PR | Ubiquinones | 1 | 2 | 3 | 4 | 2 |
|  | Polyprenols | 2 | 1 | 2 | 2 | 2 |
| SL | Diacyltrehaloses (DATs) | 3 | 1 | 1 | 1 | 1 |
|  | Sulfolipids | 2 | 0 | 3 | 1 | 4 |

The number represents the number of lipids matched by MS-LAMP in each of the isolates. S: Streptomycin; I: Isoniazid; E: Ethambutol; MDR: Multidrug resistance;
FA: Fatty Acyls; GL: Glycerolipids; GP: Glycerophospholipids; PK: Polyketides; PR: Prenol Lipids; SL: Saccharolipids;

GL group consisted of monoacylglycerols, diacylglycerols and triacylglycerols. GP consisted of glycerophosphoethanolamines (PEs), glycerophosphoglycerols (PGs), glycerophosphoinositols (PIs), glycerophosphoglycerophosphoglycerols (CLs) and glycerophosphoinositolglycans, all these in their mono- and di-acylated form. PK group consisted of linear and hybrid PKs. PR group consisted of ubiquinones and polyprenols. SL group consisted diacyltrehaloses (DATs) and sulfolipids **(Table 3)**.

Further, the lipids identified from resistant isolates were compared for similar and distinct lipids of sensitive isolate. Similar lipids were more among sensitive and IE resistant followed by MDR. More distinct lipids were among sensitive and SI resistant (51.39%), followed by isoniazid resistant (51.07%), MDR (46.78%) and IE resistant isolate (37.12%).

Among FA, I resistant isolate (25%) showed more distinct lipids followed by MDR (23.91%), SI (23.75%) and IE (16.90%) resistant isolate. Among GL, sensitive isolate showed more distinct lipids than resistant isolates. I resistant isolate had 29.35% distinct lipids, followed by IE resistant (27.72%), SI (21.54%) and MDR (19.9%) isolate. Among GP, resistant isolates showed more distinct lipids than sensitive isolate. SI resistant isolate has 51.62% distinct lipids, followed by MDR (49.71%), IE (49.08%) and I (43.72%) resistant isolate. Among PK, MDR and IE resistant isolates showed more distinct lipids than sensitive isolates, whereas SI and I resistant isolate had less distinct lipids than sensitive isolates. MDR isolate had 4.29% distinct lipids, followed by IE (3.71%), I (3.59%) and SI (1.8%) resistant isolate. Among the PR, resistant isolates showed more distinct lipids in comparison to sensitive isolate. SI isolate had 2.46 % distinct lipids, followed by IE (2.11%), MDR (1.79%) and I (1.2%) resistant isolate. Among SL, sensitive isolates had more distinct lipids than resistant isolates. SI isolate had 4.23% distinct lipids, followed by IE (2.68%), MDR (2.5%) and I (2.41%) resistant isolate.

## 4. Discussion

It is well established that Mycobacterium is rich in lipids and forms the major constituent of the cell wall. So, in the present study, lipids of first-line drug resistant and drug sensitive isolates of *M.tb* were characterised by TLC and mass spectrometry that showed significant changes in the concentrations of few of the lipids. The growth conditions can alter the lipid composition; hence organisms were grown under similar conditions (7).

### 4.1 Lipid analysis by 2D TLC

Almost all the lipids identified showed variation in resistant isolate compared to sensitive isolate, however all were statistically insignificant. Total lipid content was different among categories of resistant isolates but was statistically insignificant (Table 2).

Among the non-polar lipids, TAG was increased by ∼1.2 folds in SE resistant and R resistant isolates compared to sensitive isolates. It is established that TAGs tend to accumulate among drug resistant strains of *M.tb* (8). However, in contrast, a reduction in the TAG content though statistically insignificant was observed among SIE resistant category in our study. Though other non-polar lipids like PDIM and MQ showed variation among different isolates, it was statistically insignificant (Table 2).

Among the polar lipids, PIMs showed variations among sensitive and resistant categories of *M.tb*. AC1PIM5 was increased by ∼1.5 folds in SE resistant and reduced by ∼2.2 folds in SIE resistant isolates along with AC2PIM5 by ∼1.9 folds compared to sensitive isolates. AC2PIM2 was not detectable in R resistant and SE resistant isolates. However, correlation with monoresistant strains was not possible. PI increased significantly by ∼1.5 folds in SI resistant isolates, this correlated with monoresistant isolates of S (by ∼1.1 folds) and I (by ∼1.1 folds) which showed an increase in PI but was insignificant statistically, and this might be owed to the combination drugs (Table 2). Also, changes in the total phospholipid content of combination drug resistant isolates was reported in 1989 (9).

The results of DPG were interesting, as the combination of drugs increased the DPG levels decreased, it was absent in SIE resistant isolates, the concentration was increased in SE resistant (by ∼2.4 folds) and also in IE resistant (by ∼1.3 folds), it was still more increased in monoresistant isolates of S (by ∼1.38 folds), and I (by ∼1.56 folds) (Table 2).

Lipids identified by 2D TLC showed variation in MDR isolates, isoniazid monoresistant isolates and isoniazid-ethambutol resistant isolates but none of them was significant statistically (Table 2), where the drug targets are mainly components of cell wall. The combined effect of total lipids could be the cause which gives a characteristic feature for these isolates cell wall or some other lipids not included in this study may have a role in altering the cell wall morphology.

### 4.2 Lipid analysis by Mass spectrometry

By using 2D TLC only a few lipids could be identified, hence selected isolates were subjected to mass spectrometry analysis. The data obtained is qualitative wherein different lipids in the samples are identified but are not measured quantitatively. The MS data analysis was through positive ion mode, considering the positive adducts [(M+H)^+^ and (M+Na)^+^] as mycobacterial lipids require positive mode detection (10).

Mycolic acids (MAs) are the distinctive lipids found in Mycobacteria, MAs are targeted in antimycobacterial therapy, and they have significant immunomodulatory role during host-pathogen interactions. In the earlier studies, it has been shown that mycolic acid profile changes in response to different conditions (11, 12). In this study, all the isolates exhibited alpha-, keto- and methoxy-mycolic acids (Table 12). The overall composition is not altered, however, differences in individual MAs of the different isolates was observed. Sensitive and MDR isolate showed higher methoxy-MAs, followed by alpha- and keto-form. Whereas, in I, SI and IE resistant isolates showed increased alpha-form, but methoxy- and keto-form were in equal proportion. In a comparison of MAs between resistant and sensitive isolates, alpha- and keto-form accumulation increased in resistant isolates except for MDR. Methoxy-MAs decreased in the resistant isolates except for MDR. An *M. bovis* (BCG strain) with ethA-ethR locus mutation had lower overall MAs levels and led to the persistence of *M. bovis* in the mouse model (13). In this study, MDR isolate showed a marginal decrease in alpha- and keto-form, but a slight increase in the methoxy-MAs. The deceptive differences in MAs may contribute significatly to the susceptibility of *M.tb* to anti-TB therapies (11).

The branched fatty acids are the components of different lipids that includes PDIMs, DAT, TATs, PATs, and sulfolipids etc., (11, 14), the presence on the cell surface promote the intracellular survival of *M.tb* (15). In this study, all the samples exhibited phthioceranic acids, hydroxypthioceranic acids, mycocerosic acids and mycolipanoic acids (Table 3). Mycolipenic acid was seen only in sensitive isolate, and mycosanoic acids in all resistant isolates. These fatty acids absence could be owed to the infinitesimal quantity of these in the sample. Further confirmation is required as they are not detected by mass spectrometry or they may not be present in *M.tb* isolates.

The cell wall outer layer consists of PDIMs, sulfolipids, DATs, and PATs bound to the mycolic acids (11, 16). These are restricted in distribution and not essential for *M.tb* growth in vitro, though involved in pathogenicity (17). Present study showed insignificant increase in the total PDIMs between sensitive and resistant isolates (Table 3). DIMA was increased by 5% in the MDR isolate, but it was decreased in the other resistant isolates. In contrast, DIMB was decreased in MDR isolate and increased in others. Overall, increase in PDIMs may help *M.tb* to become impermeable to anti-TB drugs. PDIM plays an imperative role in maintaining the cell wall integrity, it is reported that PDIM mutants exhibit more permeable cell wall that was more sensitive to detergent(16). Our results are in concordance with other report, where a tesA (required for PDIM synthesis) deletion mutant of *M. marinum* revealed hyper susceptibility to antibiotics suggesting PDIMs role in the mycobacterial reaction to chemotherapeutic agents (18).

Along with major sulfolipid SL-I, SL-I’, SL-II, and SL-II’ are three minor tetra-acylated forms differing by the fatty acyl composition from SL-I. Also SL-III, a tri-acylated form was described by Goren et al., (19,20). SL contribute to early stage virulence of mycobacterial infection or glycolipids stimulation by counteracting the immunopotentiation effect of TDM (21). Present study revealed that all the isolates had relatively less number of SL (Figure 3), in that SL-I was seen among all the isolates except I resistant isolate. SL-II in SI resistant isolate and SL-III in MDR isolate. Overall, an increase in the SL was detected in resistant isolates compared to sensitive isolate (Table 3). The virulence lipids (PDIM and SLs) synthesis increases during infection (22). Hence the increased SL in the resistant isolates could be a mechanism of fitness. However, all types of SLs were undetected, as a result of infinitesimal quantity of these lipids in the samples or they may not be present in the *M.tb* isolate which needs confirmation.

TAGs occur as cell wall components and as the storage form of energy for latent *M.tb* (23). In our study, among glycerolipids, TAGs were increased in resistant isolates (49.7%) compared to sensitive isolate (45.5%) (Table 3), conconrdance with 2D-TLC observation. Interestingly, W/Beijing family strains showed TAGs accumulation under normoxic conditions, that relate to the prevalent drug resistance and high virulence (8).

Increase in total phospholipids were identified in MDR, followed by SI and IE isolate when compared to sensitive isolate (Figure 3). In a study by Sareen et al., cell wall phospholipids were shown a quantitative increase in the ethambutol resistant *M. smegmatis*, and another study by the group showed a decrease in phospholipid content among ethambutol resistant *M.tb* H37Ra (24, 25). A study by Kanwar et al., measured phospholipids quantitatively in *M. smegmatis* strain that is resistant to SE and SI where they found SI resistant isolate has fewer phospholipids than sensitive isolate (9). This is contrasting as our study is qualitative analysis and till now, no reports of phospholipids among *M.tb* clinical isolates.

Glycerophospholipids (mainly PE), LM, PIM, and ManLAM may interact with the host and are found in the cell wall outermost layers of *M.tb* and other Mycobacterium species (26). In our study, PIMs were more among the resistant isolates compared to the sensitive isolate (Table 3). Emerging data indicate vital role of PIM in the cell envelope permeability, inner membrane integrity, and cell division regulation (17). Also, PIMs have potent immunomodulatory activities important for *M.tb* pathogenesis (27).

This sort of composition changes in the lipid is also reported previously by few studies for resistant *M.tb* (9, 24) and also other bacteria (7,25, 28, 29). The lipid profile differences in the resistant and sensitive isolates do implicate a possible role of lipid composition in development of resistant phenotype in *M.tb*. The observations presented here clearly demonstrate that mycobacterial drug resistance leads to alterations in the cell wall composition. These alterations could lead to changes in the transport of the drugs across the cell or cell wall permeability.

## 5. Conclusion

To our knowledge this is the first study, that provides widespread information on the variations in the lipids among different first-line drug resistant categories such as mono-drug, bi-drug, poly-drug and multidrug resistant clinical isolates.

Owing to the fact that lipids play an imperative role in the cell structure and permeability besides potential host interaction, a production decrease or increase might not only cause resistant *M.tb*, it may extremely modify the cell surface presentation of immunomodulatory antigens and host functions interaction.

## Acknowledgments

The authors are thankful to the Director NIMHANS for permitting to carry out the study. Kind help of Neurology Faculty in case selection and performing diagnostic LP is kindly acknowledged. The authors would like to acknowledge the technical help rendered by Technicians of IISc, Bangalore for the Mass spectrometric analysis. The work was supported by University Grants Commission (UGC), New Delhi [Grant no: 20-06/2010(i)EU-IV] and Council of Scientific and Industrial Research (CSIR), New Delhi [Grant no: 09/490(0083)/2012].

